# Assessing the validity of the zero-velocity update method for sprinting speeds

**DOI:** 10.1101/2023.07.07.548103

**Authors:** Gerard Aristizábal Pla, Douglas N. Martini, Michael Potter, Wouter Hoogkamer

## Abstract

The zero-velocity update (ZUPT) method has become a popular approach to estimate foot kinematics from foot worn inertial measurement units (IMUs) during walking and running. However, the accuracy of the ZUPT method for stride parameters at sprinting speeds remains unknown, specifically when using sensors with characteristics well suited for sprinting (i.e., high accelerometer and gyroscope ranges and sampling rates). Seventeen participants performed 70-meter track sprints while wearing a Blue Trident IMeasureU IMU. Two cameras, at 20 and 70 meters from the start, were used to validate the ZUPT method on a stride-by-stride and on a cumulative distance basis. In particular, the validity of the ZUPT method was assessed for: (1) estimating a single stride length attained at the end of a 70m sprint (i.e., stride at 70m); (2) estimating cumulative distance from ∼20 to ∼70 m; and (3) estimating total distance traveled for a 70-meter track sprint. Individual stride length errors at the 70-meter mark were within - 6% to 3%, with a bias of -0.27%. Cumulative distance errors were within -4 to 2%, with biases ranging from -0.85 to -1.22%. The results of this study demonstrate the ZUPT method provides accurate estimates of stride length and cumulative distance traveled for sprinting speeds.

## Introduction

Sprinting performance is key in track and field [1] and team sports [2,3]. Sprinting performance determinants, such as maximal speed [1], can be extracted with a variety of commercially available devices such as (high speed) video cameras [4], radar guns [5], laser beams [6], smartphone apps [4], optical measurement systems [7], and resistance devices [8]. Inertial measurement units (IMUs) also have the potential to assess sprint performance. An IMU generally consists of tri-axial accelerometers, angular rate gyroscopes and magnetometers [9]. Primary benefits of IMUs are affordability, easy set-up, minimal interference with the runner’s performance, and the ability to provide instantaneous feedback.

One of the challenges of IMUs is that position measures are sensitive to integration drift [10,11]. Integration drift error can be minimized with the zero velocity update (ZUPT) method.

The IMU-based ZUPT method can provide estimates of foot kinematics with shoe-worn IMUs in real-life settings [12,13,14,15]. The ZUPT method works in a four-step process. In the first step, the stride segmentation, raw IMU signals are used to detect zero velocity instants when the foot is (approximately) stationary on the ground. In the second step, the rotational orientation estimation, IMU orientation in space is estimated using an extended Kalman filter. In the third step, the translational velocity estimation, linear accelerations are integrated between two successive zero velocity instants. Exploiting the fact that the foot is at zero velocity at the start and end of each stride, a linear drift correction is performed to correct the velocity back to zero during each zero-velocity instant, thus reducing the integration drift errors. In the fourth step, the trajectory formation, stride parameters (e.g., stride length) are obtained by integrating foot velocities. While the ZUPT method has high validity in walking [13] due to clear zero-velocity foot periods, external validity of the ZUPT method during running and sprinting could be limited as the foot may not have a clear zero-velocity period. In addition, peak accelerations and angular velocities during sprinting might be outside the measurement range of commercially available IMUs, potentially further impacting the external validity of stride lengths obtained with the ZUPT method.

The traditional ZUPT method [10] has been shown to yield accurate estimates at running speeds up to 6.4m/s [12,16,17,18]. Bailey and Harle [16] investigated the accuracy of the traditional ZUPT method for speeds up to 3.4m/s on a treadmill. Brahms et al. [18] investigated the accuracy of the traditional ZUPT method for speeds up to 4.36m/s during overground running. Bailey and Harle [17] and Potter et al. [12] additionally investigated the effects of several IMU specifications. Bailey and Harle [17] investigated the effect of accelerometer range and sampling frequency, while Potter et al. [12] investigated the effect of accelerometer range, gyroscope range and sampling frequency. The study by Potter et al. [12] was conducted in outdoor environments and the speeds analyzed were much higher (up to 6.4m/s) than those analyzed by Bailey and Harle [17] (up to 3.4m/s). Therefore, the ZUPT estimates obtained by Potter et al. [12] were more impacted by limitations in the IMU hardware (i.e., specifications). Quantifying stride metrics with the traditional ZUPT method at speeds up to 6.4 m/s over 100 meters showed that sampling frequency significantly impacts the traditional ZUPT estimates of individual stride parameters [12]. In addition, saturation due to low gyroscope range (i.e., 750 °/s) or accelerometer range (i.e., 24g) also reduced stride estimate accuracy [12]. Though, the authors noted that acceptable estimates (i.e., errors below 5%) may still be obtained if the amount of data that is lost due to saturation remains small (i.e., 1.5% in acceleration signals and 2.6% in angular velocity signals) 74 [12].

De Ruiter et al. [14] assessed the validity of the ZUPT method for sprinting speeds (i.e., speeds over 8m/s), using an IMU with a ±16g and a ±2000°/s range, sampling at 500Hz. To mitigate the effects of accelerometer saturation, the ZUPT method was modified with respect to the traditional ZUPT method [14]. De Ruiter et al. [14] defined a stride by the time interval between two consecutive initial contacts rather than between two consecutive zero-velocity times. Rather than applying a linear drift correction, integration drift error in velocity was corrected by first identifying the minimum values (i.e., velocity offsets) in the filtered velocity signals between 20 and 100 milliseconds following initial contact, then subtraction of velocity offsets from the raw velocity signals and imposing all data points prior velocity offsets to be zero [14]. Finally, offset corrected velocities were integrated to obtain stride length. For peak sprint speeds of 8.42±0.85m/s, the obtained bias and limits of agreement were reasonably accurate (i.e., -2.51±8.54%) [14]. Based on the findings by Potter et al. [10], the sprint speed results reported by De Ruiter et al. [14] could have been affected by the IMU specifications, especially their relatively low accelerometer range (±16g). This suggests that the accuracy of the traditional ZUPT method on a stride-by-stride basis for track sprinting speeds could be improved when using sensors with characteristics better suited for sprinting (i.e., higher ranges and sampling rates).

Therefore, the aim of this study was to assess the validity of the traditional ZUPT method for: (1) estimating a single stride attained at the end of a 70m sprint (i.e., stride at 70m); (2) estimating cumulative distance from ∼20 to ∼70 m; and (3) estimating total distance traveled for a 70-meter track sprint. These estimations were performed using data collected with an IMU with a high accelerometer range (±200g) and sampling frequency (1600Hz). Based on Potter et al. [12], we hypothesized that IMU-based stride lengths and cumulative distances would be within 5% from stride estimates obtained with a camera-based capture system.

## Methods

### Participants

Seventeen participants over 18 years old were enrolled in this study (Table 1; recruitment period: March 14, 2022 – May 25, 2022). Inclusion criteria were ability to run 7m/s or faster and being free of injury for at least three months prior their testing session. Exclusion criteria included any orthopedic, cardiovascular, or neuromuscular conditions that would affect sprint performance. Each participant provided a written informed consent approved by the University of Massachusetts Amherst Institutional Review Board (#3143).

**Table 1.**
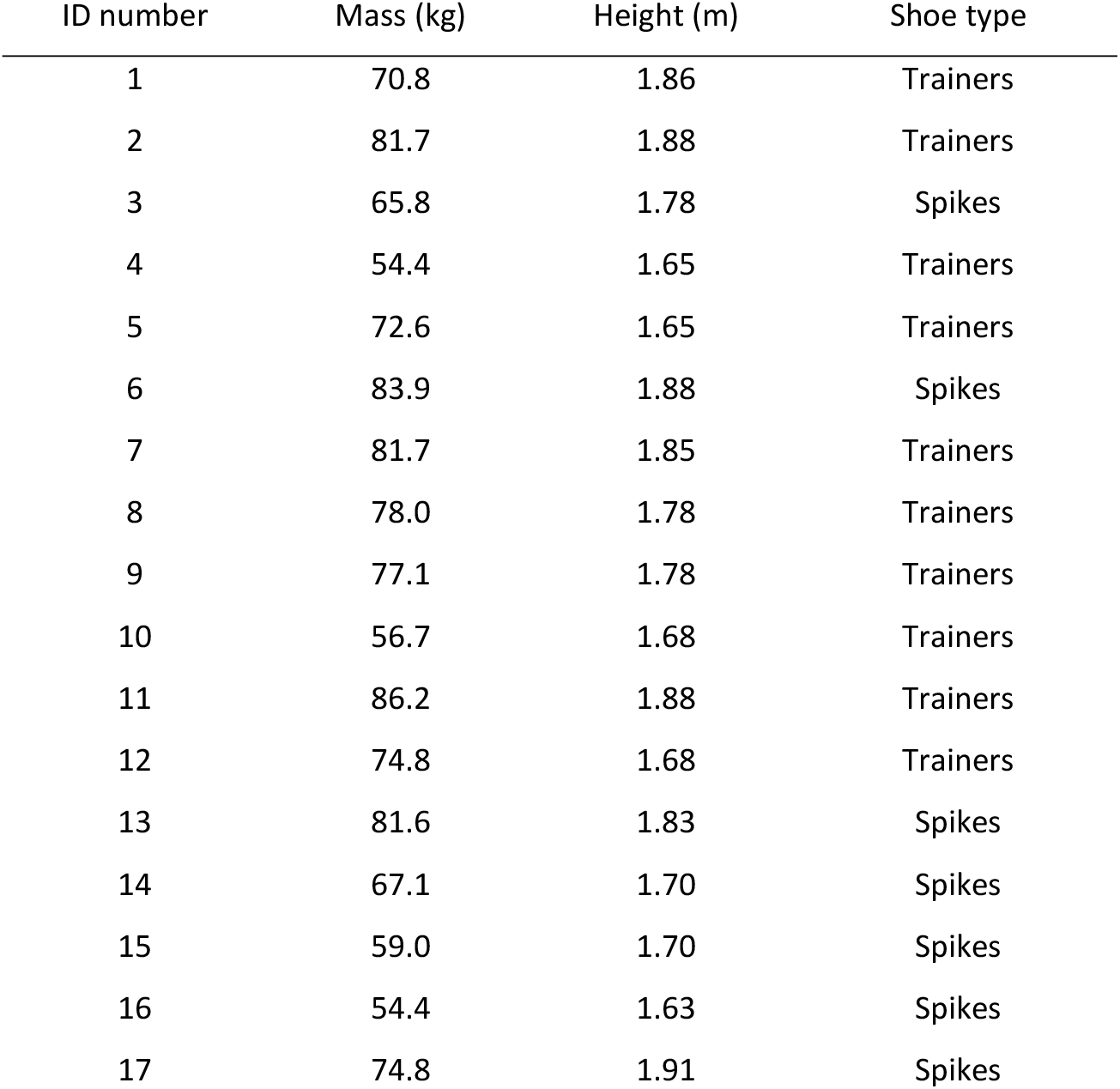
Descriptive characteristics for all participants.

| ID number | Mass (kg) | Height (m) | Shoe type |
| --- | --- | --- | --- |
| 1 | 70.8 | 1.86 | Trainers |
| 2 | 81.7 | 1.88 | Trainers |
| 3 | 65.8 | 1.78 | Spikes |
| 4 | 54.4 | 1.65 | Trainers |
| 5 | 72.6 | 1.65 | Trainers |
| 6 | 83.9 | 1.88 | Spikes |
| 7 | 81.7 | 1.85 | Trainers |
| 8 | 78.0 | 1.78 | Trainers |
| 9 | 77.1 | 1.78 | Trainers |
| 10 | 56.7 | 1.68 | Trainers |
| 11 | 86.2 | 1.88 | Trainers |
| 12 | 74.8 | 1.68 | Trainers |
| 13 | 81.6 | 1.83 | Spikes |
| 14 | 67.1 | 1.70 | Spikes |
| 15 | 59.0 | 1.70 | Spikes |
| 16 | 54.4 | 1.63 | Spikes |
| 17 | 74.8 | 1.91 | Spikes |

### Experimental protocol

We collected IMU data using the commercially available Blue Trident IMeasureU IMU (see Table 2 for sensor characteristics). We placed the IMU on the right shoe (Fig 1); specifically, to the medial dorsal aspect of the foot [12]. The IMU was attached using double-sided tape and strapped down with Hypafix tape to reduce motion artefacts.

**Table 2.** Blue Trident IMeasureU IMU sensor specifications.

|  |  |
| --- | --- |
| Accelerometer sample frequency (Hz) | 1125, 1600 |
| Accelerometer range (g) | $\pm 16, \pm 200$ |
| Accelerometer bit-resolution (bits) | 16, 13 |
| Accelerometer resolution ($\text{m/s}^2$) | 0.0048, 0.48 |
| Accelerometer bandwidth (Hz) | 1125, 1600 |
| Accelerometer anti-aliasing filter cut-off frequency (Hz) | 473, 800 |
| Accelerometer noise density ($\text{mg}/\sqrt{\text{Hz}}$) | 0.23 |
| Gyroscope sample frequency (Hz) | 1125 |
| Gyroscope range ($\text{deg/s}$) | $\pm 2000$ |
| Gyroscope bit-resolution (bits) | 16 |
| Gyroscope resolution ($\text{deg/s}$) | 0.061 |
| Gyroscope bandwidth (Hz) | 773.5 |
| Gyroscope anti-aliasing filter cut-off frequency (Hz) | 473 |
| Gyroscope noise density ($\text{deg/s}/\sqrt{\text{Hz}}$) | 0.015 |

*Left values correspond to the low-g accelerometer. Right values correspond to the high-g accelerometer.*

**Fig 1.**
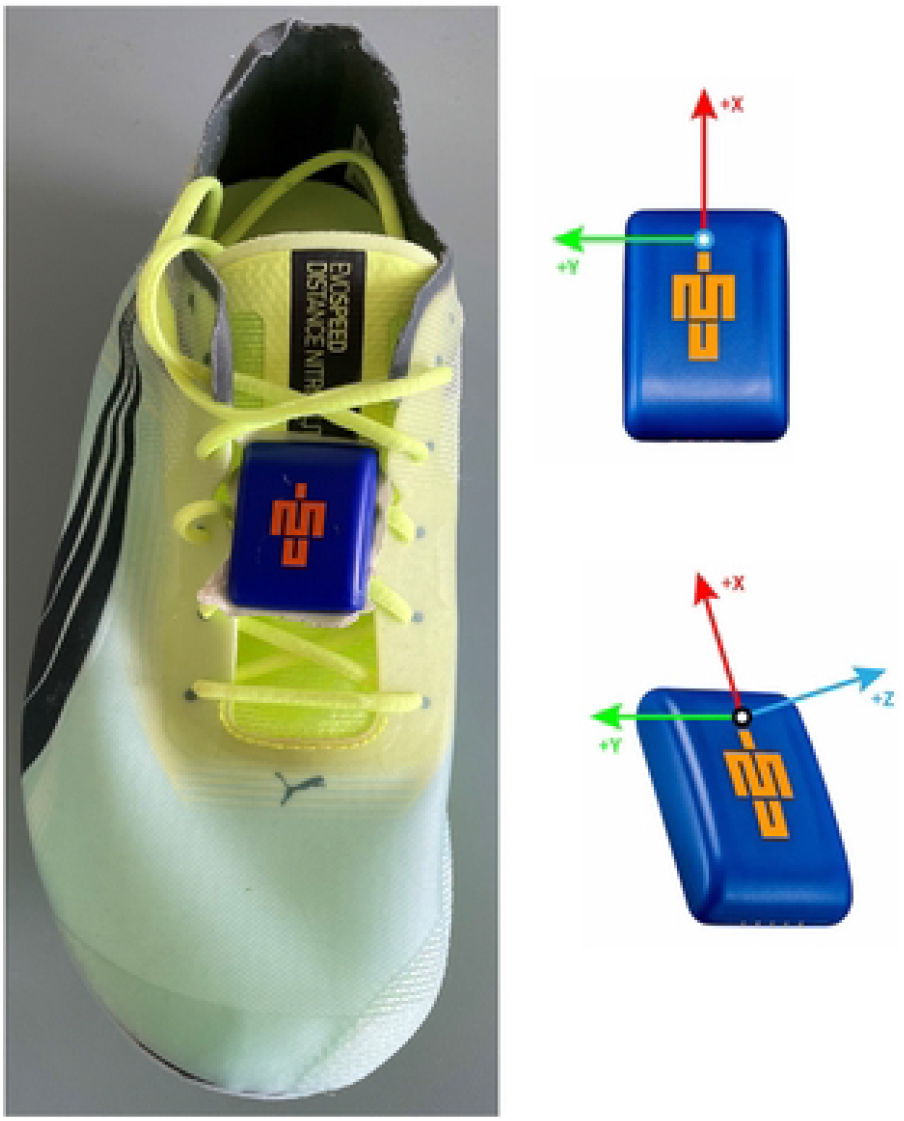
Blue Trident IMeasureU attachment to the instep of the shoe.

We placed two cameras (Apple iPhone 12, 1080 pixels at 240Hz), mounted on tripods to capture footfalls at 20m and 70m to establish the exact distance traveled from the start to each footfall closest to each of these marks. Furthermore, the camera at the 70m mark captured the distance between two consecutive footfalls for the right foot to validate the traditional IMU-based ZUPT method for a single stride. Following de Ruiter et al. [14], we placed the iPhone cameras 8 meters away from the track lane that subjects were running in. We choose to place the cameras at 20 and 70m to ensure participants were running near top speed (at 70m) and to separate the acceleration phase from the maximal speed phase (at 20m). We placed tape marks at set distances throughout the full capture space to calibrate the iPhone camera views and to account for triangulation in the analysis (see below). The distance between consecutive tape marks was seven meters. We used an additional mobile camera at 40m to count the number of strides taken until the participants ran through the view of the static cameras. This enabled us to compare the same stride from the IMU and each stationary iPhone camera.

Each subject completed a self-selected warm-up, then ran a 70-meter sprint at maximal effort on an outdoor track. We instructed subjects to stand still for ∼15 seconds before the start of the sprint with the vertical projection of the IMU aligned with the start line. This was done to subtract the gyroscope fixed bias [19].

### IMU analysis

The raw IMU signals were downloaded and analyzed with customized software in Python (Python Software Foundation, Delaware, DE, USA). The customized software used the high-g accelerometer (±200g) whenever the low-g accelerometer (±16g) saturated. We included this step because the resolution of the high-g accelerometer is much lower than the resolution of the low-g accelerometer. Thus, it would not be possible to obtain accurate IMU-derived stride lengths by just using the high-g accelerometer. Then, we calculated stride lengths using the ZUPT method, introduced above and described in detail in Potter et al. [10]. The detection of stationary periods (i.e., time points with minimum angular velocity during detected stance phases) was done using customized software in Python (Python Software Foundation, Delaware, DE, USA).

### Validation

Individual stride lengths were added to yield the estimated total distance traveled (D_calc_). We calculated the cumulative distance error (D_err_), for 0-∼20, ∼20-∼70 and 0-∼70 meters, as follows [10]:

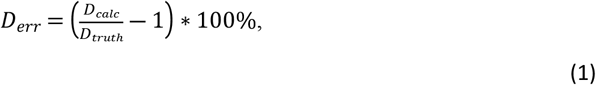

where D_truth_ is the known distance traveled (i.e., ∼20, ∼50 and ∼70 meters), obtained from the camera recordings. Total known distances traveled were divided by the duration extracted from the IMU to obtain average speeds.

Individual IMU-based stride lengths errors (S_err_) were calculated for the stride nearest to the 70-meter mark as follows:

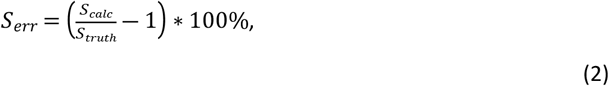

where S_truth_ is the known stride length obtained from the camera and S_calc_ is the stride length obtained with the IMU-based ZUPT method. The stride length was divided by the IMU-derived stride duration (i.e., the time between two consecutive stationary periods) to obtain stride speed.

S_truth_ and D_truth_ were calculated by tracking the IMU’s position using Kinovea software (www.kinovea.org). We used known points (i.e., tape marks) and the known distances between those points to define a perspective grid. That was done to obtain a calibrated camera view that allowed for accurate calculation of stride lengths and the exact total distance traveled.

Three trained researchers independently analyzed all the videos with Kinovea. If the difference between the results obtained from the three researchers for the same trial was smaller than 2%, then the average of the three was taken. If the difference was larger than 2%, we removed the results from the researcher whose result was further away and the average of the two remaining researchers was taken. If the average of the two still did not lead to a difference smaller than 2%, then all researchers reanalyzed the same trial and the process was repeated.

### Statistical analysis

We used Bland-Altman analysis to determine the agreement between the individual stride and cumulative distances obtained with Kinovea and with the IMU. We used simple and multiple linear regressions to investigate the effect of body mass and speed on the individual stride and cumulative distance errors. For the simple linear regressions, errors were included as dependent variables and the speeds were independent variables. For the multiple linear regressions, errors were included as dependent variables and the speeds and masses as independent variables. Alpha level was set *a priori* to 0.05 for all analyses.

## Results

### Effect of sprinting speeds and body mass on stride length estimation

Participants reached stride speeds ranging from 6.0m/s to 9.3m/s (8.00±0.88m/s). We found that the **i**ndividual stride length errors (Fig 2A) were between -6 and 3% for the individual stride at 70m, with a bias (± limits of agreement) of -0.27 ± 4.61% (Fig 3A). Participants reached 20-meter with average speeds ranging from 4.5m/s to 6.3m/s (5.51±0.51m/s). We found that 20-meter cumulative distance errors (Fig 2B) were smaller, ranging from -4 to 2%, with a bias of -0.85 ± 3.47%) (Fig 3B). Participants ran 20-70-meter with average speeds ranging from 6.7m/s to 9.5m/s (8.29±0.78m/s), while 20–70-meter cumulative distance errors (Fig 2C) remained within -4 to 2%, with a bias of -1.22 ± 3.67% (Fig 3C). Participants ran the full 70-meter sprints with average speeds ranging from 6.0m/s to 8.2m/s (7.19±0.65m/s), with 70-meter cumulative distance errors (Fig 2D) within -4 to 2%, with a bias of -1.11 ± 3.50% (Fig 3D). Note that average speeds for 20-70m (Fig 2C) were higher than those for the single stride at 70m (Fig 2A; Fig 4). Individual and multiple linear regression for the stride and cumulative distance errors are presented in Table 3. Speed and body mass had a significant effect on the individual IMU-based stride length error. Speed and body mass had a significant effect on the 20-70-meter cumulative distance error. Speed had a significant effect on the 70-meter cumulative distance error.

**Table 3.** Simple and multiple linear regression results. Significant differences are highlighted in bold.

| Error | Linear regression type | *p* value | Adjusted R<sup>2</sup> value | Standardized $\beta$ speed | Standardized $\beta$ mass |
| --- | --- | --- | --- | --- | --- |
| Stride 70m | Simple | **0.01** | 0.32 | -0.6 |  |
|  | Multiple | **0.02** | 0.34 | -0.68 | 0.26 |
| Cumulative 20m | Simple | 0.52 | -0.037 | -0.17 |  |
|  | Multiple | 0.79 | -0.011 | -0.20 | 0.077 |
| Cumulative 20-70m | Simple | **0.0044** | 0.39 | -0.65 |  |
|  | Multiple | **0.0093** | 0.41 | -0.74 | 0.26 |
| Cumulative 70 | Simple | **0.037** | 0.21 | -0.51 |  |
|  | Multiple | 0.079 | 0.18 | -0.60 | 0.23 |

**Fig 2.**
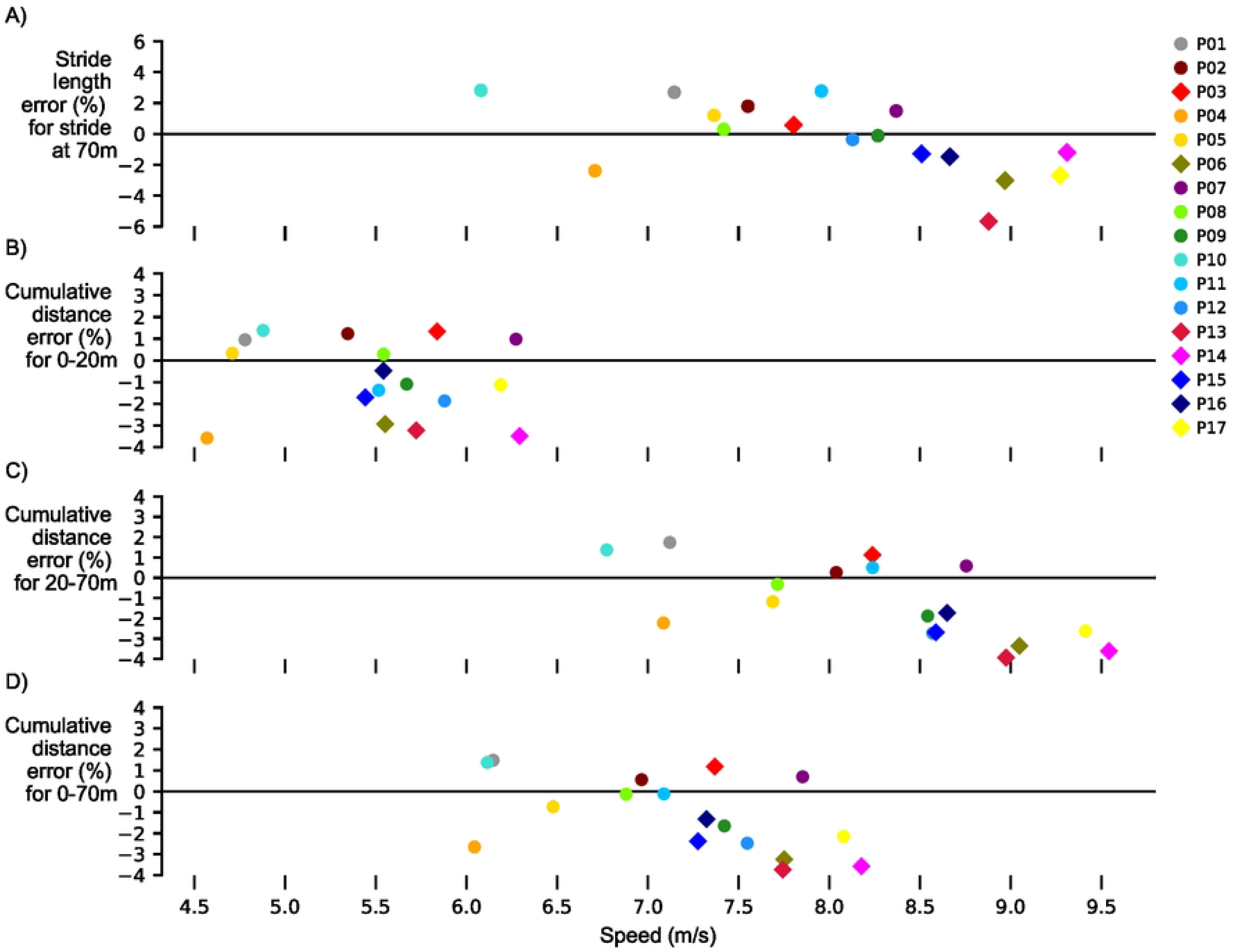
Stride length error and cumulative distances errors versus average speed comparing IMU to Kinovea. Stride length errors were between -6 and 4% across speeds, while cumulative distance errors were between -4 and 2% across speeds. Colors represent different subjects. Circles represent participants wearing trainers and diamonds represent participants wearing spikes.

**Fig 3.**
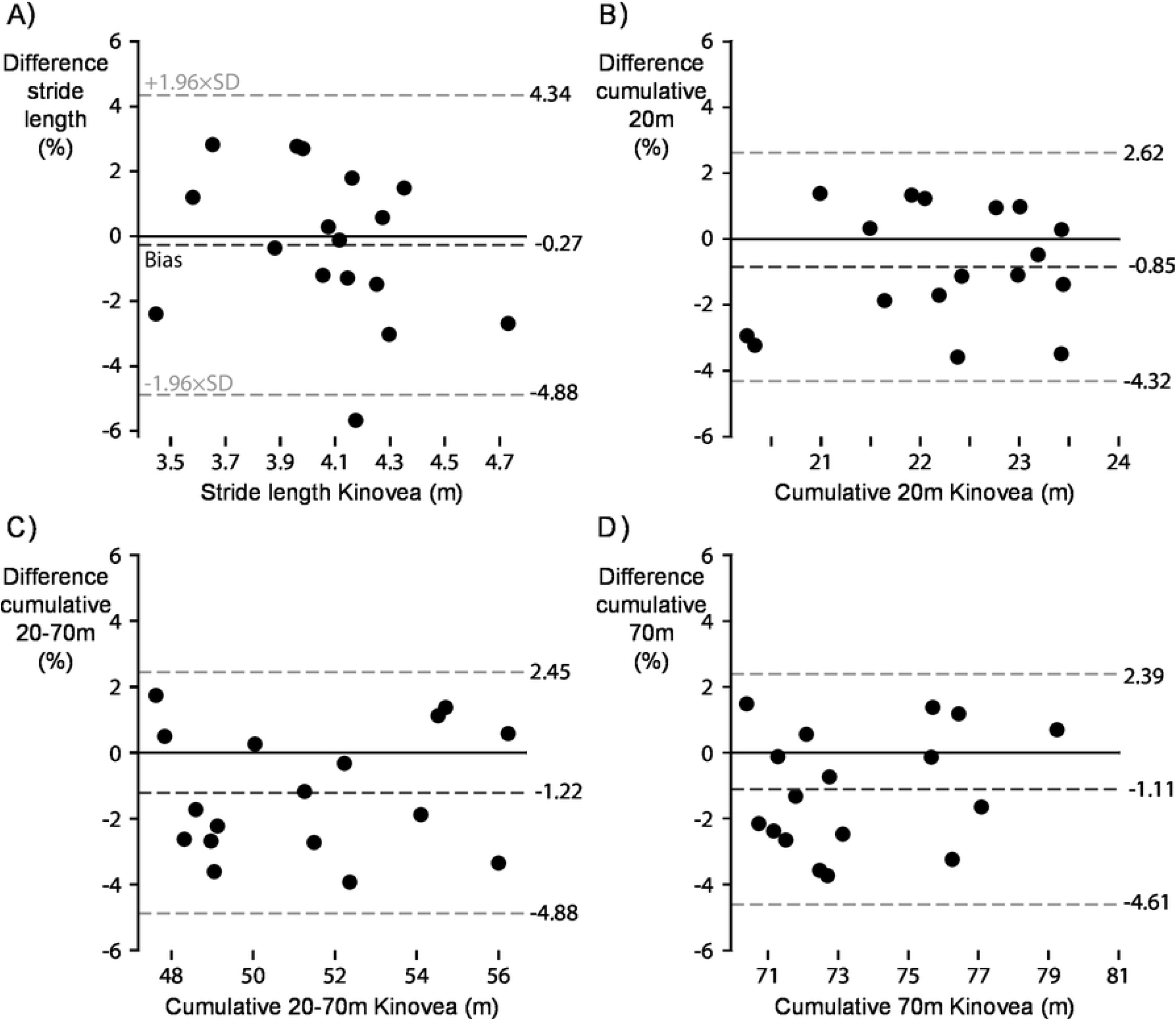
Comparison of IMU-derived measures against Kinovea-derived measures. **A** For the individual stride length at 70m the average bias (dashed line black) ± limits of agreement (dashed line grey) were - 0.27 ± 4.61%. **B** For cumulative distance over the first 20m the average bias ± limits of agreement were - 0.85 ± 3.47%. **C** For cumulative distance from 20 to 70m the average bias ± limits of agreement were - 1.22 ± 3.67%. **D** For cumulative distance over the full 70m the average bias ± limits of agreement were - 207 1.11 ± 3.50%.

**Fig 4.**
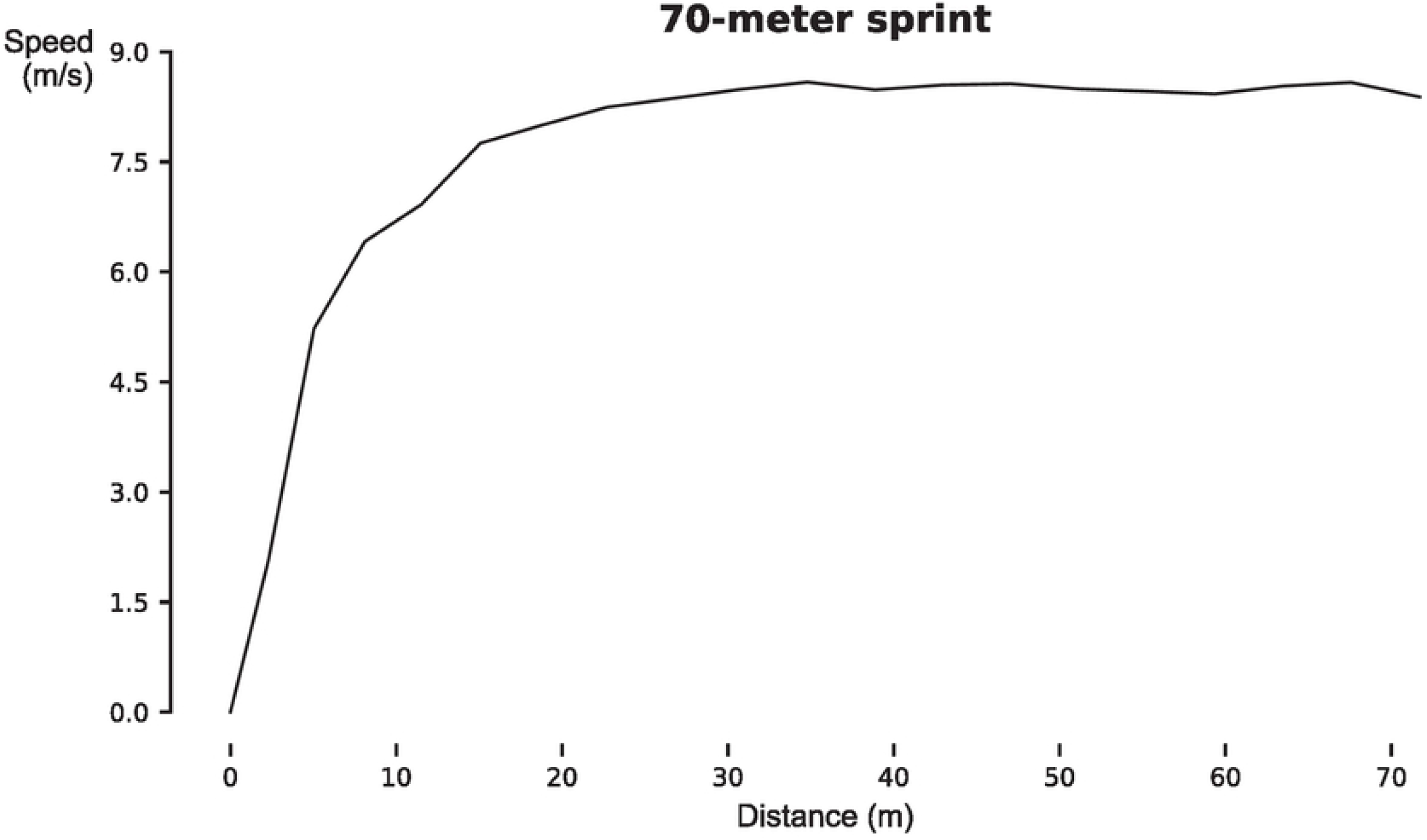
IMU-derived speed for a 70-meter sprint on a stride-by-stride basis as a function of distance traveled for a representative subject (P09).

## Discussion

The aim of this study was to evaluate the accuracy of the IMU-based ZUPT method on a stride-by-stride basis and on total distance traveled for track sprinting. We found that individual stride and cumulative distance errors were low (i.e., within -6 to 3% and -4 to 2%, respectively).

For peak sprinting speeds of 8.00 ± 0.88 m/s, we obtained an average bias ± limits of agreement of - 0.27 ± 4.61% (Fig 2). Our findings build further on limited available literature [14], expanding the validity of the IMU-based ZUPT method for sprinting speeds. For peak sprinting speeds of 8.42 ± 0.85 m/s, de Ruiter et al. [14] obtained an average bias ± limits of agreement of -2.51 ± 8.54%. To mitigate accelerometer saturation, the de Ruiter et al. provided an alternative approach of the traditional ZUPT method [10]. However, their results were likely still affected by accelerometer saturation, especially during ground contact, as suggested by evidence that accelerometer saturation leads to a degradation in accuracy of the ZUPT method [12]. The present study demonstrates that without the presence of accelerometer saturation, the traditional ZUPT method provides accurate and superior estimates for sprinting speeds over the modified ZUPT method.

Without accelerometer saturation, we obtained stride length errors within -6 to 3% for stride speeds up to 9.3m/s. Potter et al. found that for 100-meter average speeds of 6.5m/s, a ±16g accelerometer led to a 15% underestimation of a 100-meter distance, compared to a 100g accelerometer [12]. De Ruiter et al. [14] obtained stride length errors as high as 30% for stride speeds of 9m/s using a ±16g range IMU.

Our results indicate that the accuracy of the ZUPT method on a stride-by-stride basis for high sprinting speeds can be improved when using sensors with specifications better suited for sprinting velocities (i.e., higher accelerometer ranges). In addition, we observed 20-70-meter cumulative distance errors within - 4 to 2%, demonstrating that the ZUPT method yields acceptable results (i.e., errors of 5% or less) for stride speeds as high as 9.5m/s.

One of the main potential sources of error of the ZUPT method could have been gyroscope saturation during ground contact. Similarly to Potter et al. [12], we found that gyroscope saturation led to underestimated stride lengths and cumulative distance errors. We observed that underestimation increased with running speed and for participants wearing spikes. Participants wearing spikes exhibited gyroscope saturation in more steps than participants wearing trainers, which could lead to a larger percent of gyroscope data lost due to saturation. This saturation was mostly exhibited at ground contact when the foot was in plantarflexion. Trainers have thicker midsole foams than spikes [20], which provide more cushioning, likely acting as a physical low pass filter that reduces peak magnitudes in kinematics. For sprinting in spikes with less cushioning than trainers, higher peak angular velocities can be expected, which could lead to gyroscope saturation at high sprinting speeds. Potter et al. [12] found that the cumulative distance errors remained below 5% when the percentage of gyroscope data lost due to saturation was below 2.6% [12]. This suggests that the amount of data that we lost from saturation was low, even though we could not quantify what percentage of data we lost due to saturation.

For the participants that did not exhibit gyroscope saturation (n = 6), we obtained cumulative and individual distance errors within -3 to 3% for speeds up to 8.5m/s. Those errors are very similar to error reported for level walking [10]. Though we did not observe any signal saturation for these participants, that does not mean that there was no saturation, as theoretically signal saturation could be camouflaged by IMU internal preprocessing. However, the internal preprocessing of the Blue Trident IMeasureU consists of a digital low band pass filter with a cut-off frequency of 473Hz, while the sensor bandwidth is 773.5Hz. Those cut-off frequencies are well beyond what can be expected during human sprinting.

Without the presence of IMU saturation, the accuracy of estimated stride lengths during track sprinting could be of practical relevance. Accurate stride length estimations allow for accurate stride speed calculation. From the sprint speed curve presented in Fig 4, sprinting performance determinants such as maximal speed [1] can be extracted. In addition to track and field sports [1], sprinting performance is key in team sports [2, 3]. Thus, this method could provide coaches with key determinants to assess and improve sports performance. Note that this method only requires a single IMU attached to the athlete’s foot and does not require any information about IMU orientation in space and is minimally affected by subject-specific characteristics (i.e., body mass). Note that the significant effect in Table 3 of body mass could have been confounded by other factors. Future research should try to assess the validity of the ZUPT method for high sprinting speeds (i.e., speeds over 9 m/s) with sensors that have specifications better suited for sprinting. Such sensors would not admit gyroscope saturation (>±2400 °/s) and have good resolutions (>16bits) and appropriate bandwidths (>500 Hz) and sample frequencies (>1000 Hz) to capture high frequency impacts during ground contact. Future research should directly compare the effects of spikes and trainers on the accuracy of the ZUPT method, as our interpretation on the effects of spikes could have been confounded by the fact that spikes were worn for different subjects. Our findings suggest that with the appropriate sensors, the ZUPT method could possibly be used to compare the performance of different track spikes [20] as well as to test elite sprinters.

In conclusion, the results of this study demonstrate the accuracy of the ZUPT method for sprinting speeds, even with the presence of gyroscope saturation. Cumulative and individual distance errors remained within -6 to 3% for speeds ranging from 6m/s to 9.5m/s.

## Acknowledgments

The authors would like to thank UMILL lab members who helped with data collections, Shane Schwartz and Herlandt Lino for their help provided with the Kinovea analyses, Dr. Alex Shorter and Loubna Baroudi for sharing their insights and Dr. Leia Stirling for her help provided with software development and ZUPT implementation.

## Notes

### Competing Interest Statement

The authors have declared no competing interest.

